# Changes in extracellular matrix cause RPE cells to make basal deposits and activate the alternative complement pathway

**DOI:** 10.1101/152835

**Authors:** Rosario Fernandez-Godino, Kinga M. Bujakowska, Eric A. Pierce

## Abstract

The design of efficient therapies for age-related macular degeneration (AMD) is limited by our understanding of the pathogenesis of basal deposits, which form between retinal pigment epithelium (RPE) and Bruch’s membrane (BrM) early in disease, and involve activation of the complement system. To investigate the roles of BrM, RPE and complement in AMD, we generated ARPE-19 cells with the p.R345W mutation in *EFEMP1*, which causes early-onset macular degeneration. The ARPE-19-*EFEMP1*^R345W/R345W^ cells make abnormal extracellular matrix (ECM) that binds active complement C3 and causes the formation of basal deposits by normal human fetal (hf)RPE cells. hfRPE cells grown on abnormal ECM or BrM explants from AMD donors show chronic activation of the alternative complement pathway by excessive deposition of C3b. This process is exacerbated by impaired ECM turnover via increased matrix metalloproteinase-2 (MMP-2) activity. Therapies that target ECM synthesis and turnover and activation of C3 could be effective for early AMD.

## INTRODUCTION

Age-related macular degeneration (AMD) is the most common cause of blindness among elderly people, but the design of additional therapies for AMD is limited by the lack of knowledge regarding the mechanisms associated with early steps in disease [1]. Currently available treatments target intermediate/late stages of disease. For example, antioxidant supplementation can slow AMD progression in some patients with intermediate disease [2], and the development of anti-VEGF therapies has been a major advance in the treatment of choroidal neovascularization associated with late stage disease [1].However, there are no effective therapies available to prevent progression of early AMD, when basal deposits and drusen form between the basal lamina of the retinal pigment epithelium (RPE) and Bruch’s membrane (BrM), to the later vision-threatening stages of disease.

Aging causes degeneration and thickening of BrM by accumulation of extracellular matrix (ECM) structural components and impaired matrix metalloproteinase (MMP) activity, which is thought to be exacerbated by both genetic and environmental factors in AMD [3-7]. With age, RPE function will likely be affected by increased deposition and abnormal distribution of the ECM. Importantly, RPE cells synthesize MMP-2, which is key for ECM remodeling in BrM [8, 9], and dysregulation of which triggers inflammatory processes associated with the development of AMD, likely via cleavage and regulation of chemokines, ECM proteins and their regulators[4, 6, 10-12]. But identification of the specific factors and mechanisms that lead from basal deposits to drusen formation and then to progression of AMD remain to be defined.

Experimental and clinical evidence strongly indicate a role for the complement system in AMD as early as drusen formation[13-19]. Nonetheless, several anti-complement drugs that target specific complement pathways have been developed and tested in clinical trials without significant success to date [20-22]. For example, Compstatin, a peptide that inhibits activation of C3-convertase did not show efficacy in Phase II Clinical Trials [23, 24] (Clin Trial NCT01603043). This is a noteworthy result because activation of C3 is a common endpoint of all three complement pathways (classical, lectin and alternative) [25].

The design of effective complement-modulating therapies for AMD requires a better understanding of direct functional connections between alterations in the complement system and the pathogenesis of macular degeneration. Also, there is controversy about which pathway triggers complement activation in AMD and if this process is local or systemic [14, 26]. We previously demonstrated that abnormalities in the ECM cause local activation of complement by the RPE in a mouse model of macular degeneration [27-29]. Indeed, most complement risk alleles as well as complement components found in the BrM and drusen of AMD patients belong to the alternative pathway [15, 16, 18]. Under normal conditions, the complement system is constantly activated by the tick-over process at a low level. In this way, C3 is cleaved by physiological hydrolysis of its internal thioester, generating free C3(H2O) and C3a [30-32]. C3(H2O) deposited on the ECM can form a stable properdin-independent C3(H2O)-convertase capable of activating the alternative pathway, generating more C3b, which in turn creates a positive feedback loop that will result in the increased release of the anaphylatoxin C3a [33-35]. This process cannot be regulated by CFH [31, 36, 37] or inactivated by Compstatin, which may explain the inefficiency of this drug in AMD patients [23].

To further investigate how alterations in the ECM increase activation of the complement system in early macular degeneration, we generated human ARPE-19 cells with the pathogenic p.R345W mutation in the *EFEMP1* gene, and studied the response of normal human fetal (hf) RPE cells to the abnormal ECM made by the mutant ARPE-19 cells. We also investigated the response of normal hfRPE cells to BrM from eyes with AMD. The data obtained from these studies show that abnormalities in the structure and composition of the ECM, caused either by the p.R345W mutation in EFEMP1 or associated with AMD, are sufficient to produce increased complement activation and basal deposit formation by normal RPE cells. The data further suggest that C3 produced by RPE cells is activated likely via tick-over and deposited in excess on abnormal ECM, where it causes a local chronic activation of the alternative complement pathway. To our knowledge, this is the first demonstration that abnormal matrix can initiate the local activation of the complement system as one of the early steps in the pathogenesis of AMD, and that this mechanism is shared between an inherited macular degeneration and AMD.

## RESULTS

### 1. Generation of ARPE-19 cells that harbor the mutation c.1033C>T (p.R345W) in the EFEMP1 gene via CRISPR-Cas9 editing.

We have previously demonstrated that primary mouse RPE cells carrying the mutation p.R345W (c.1033C>T) in the *Efemp1* gene make basal deposits *in vitro* [29]. Given that *EFEMP1*-associated macular degeneration is an ultra-rare disease, it is difficult to obtain samples from affected patients to use for studies of basal deposit formation *in vitro* by mutant human RPE cells. However, genome editing using the Clustered regularly interspaced short palindromic repeats (CRISPR)-Cas9 system allows the assessment of functional consequences of single risk alleles on the cellular phenotype in an isogenic background [38]. Hence, we engineered ARPE-19 cells to be homozygous for c.1033C>T mutation in exon 9 of the *EFEMP1* gene (Fig 1A).

**Figure 1.**
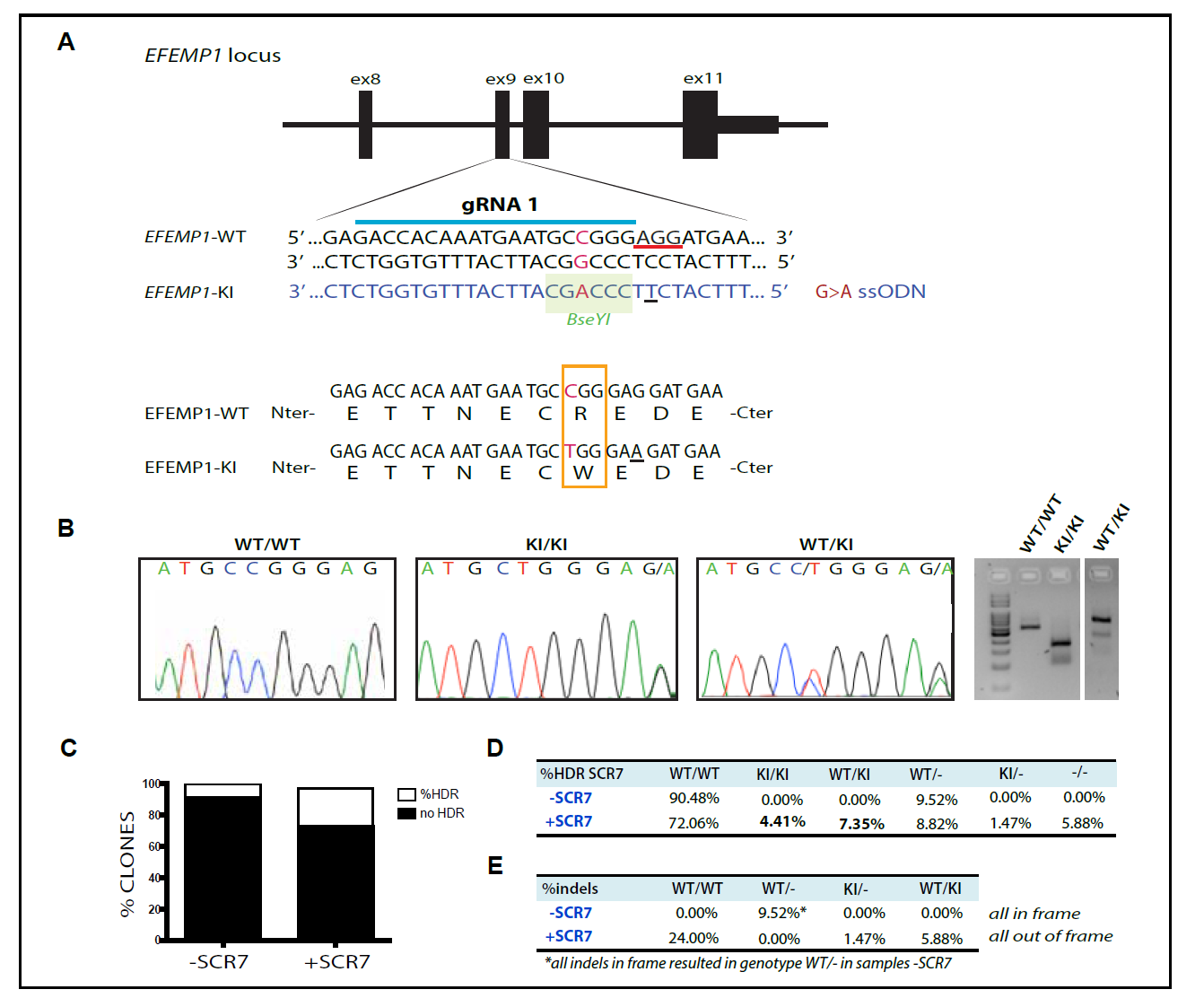
**(A)** Design of guide RNA (gRNA) and single strand oligonucleotide donor (ssODN) to knock-in the mutation c.1033C>T in the exon 9 of the *EFEMP1* gene in ARPE-19 cells via CRISPR-Cas9.PAM sequence underlined in red. Change G>A in the ssODN creates a new restriction site for the *BseYI* enzyme (highlighted in green). **(B)** Sanger sequence and agarose gel of successfully edited and isolated clones with genotype WT/WT (wildtype), KI/KI (two copies of the mutation, one per allele) and WT/KI (only one allele has the mutation). **(C)** Percentage of clones that had the mutation, comparing untreated (-SCR7) and treated (+SCR7) with the DNA ligase IV inhibitor SCR7. **(D)** Quantification of edited clones according to their genotype: WT/WT: both alleles wildtype, KI/KI: both alleles mutant, WT/KI: one allele mutant and one WT, WT/-: one allele wildtype and one allele with indels that result in null allele, KI/-: one allele with the mutation and one allele with indels that result in null allele, -/-: both alleles have indels that result in null alleles. **(E)** Percentage of clones treated with or without SCR7 which alleles had indels in frame or out of frame.

Genome editing with wildtype CRISPR-Cas9 relies on a double strand break (DSB) introduced by the Cas9 nuclease. The ability to precisely target a specific site is critical, because point mutations are generated via homology directed repair (HDR) with a single stranded donor oligonucleotide (ssODN), which carries the desired change [38, 39]. We generated a DSB after the nucleotide base 1033, which coincides with the site of mutation. We co-transfected the cells with the pSpCas9(BB)-2A-GFP (PX458) plasmid carrying the Cas9 nuclease and guide RNA (gRNA) with the ssODN harboring the desired mutation c.1033C>T. A silent mutation was introduced to change the PAM sequence of the ssODN, so the targeted allele cannot be recognized by the nuclease after HDR (Fig. 1A, B).

A total of 2208 transfected cells were plated into 96-well plates, and 937 clones derived from single cells were expanded for further analyses. In general, the efficiency of HDR using CRISPR/Cas9 is quite low due to the preferential non-homologous end joining (NHEJ) of DNA ends. However, treatment with the ligase IV inhibitor SCR7 increased the efficiency of the HDR over the NHEJ resulting in more than 25% of the total clones screened carrying the c.1033C>T mutation in one or two alleles (Fig. 1C). More than 11% of clones had at least one copy of the mutation with no evidence off-target variation, and 4.4% of them had two copies of the mutation after a single round of transfection (Fig. 1C, D). In some cases, HDR was successful but either one or both alleles also included a small indel (usually 1-8 bases) that generated hemizygous *EFEMP1*^WT/-^(WT/-), *EFEMP1*R^345W/^-(KI/-), or knock-out *EFEMP1*^-/-^(-/-) cells. Hemizygous KI/-that harbor the mutation in one or both alleles plus indels in one allele were only found in the samples treated with SCR7 (Fig. 1D). Moreover, the treatment with SCR7 appeared to favor the formation of out-of-frame indels at the expense of in-frame indels (Fig. 1E).

The presence of predicted potential off-target alterations was ruled out by Sanger sequencing (see Methods section). Only one clone presented an intronic point mutation A>G (chr13:+71022300) that did not affect the cDNA sequence. This clone was not used for further experiments.

### 2. CRISPR-Cas9 edited ARPE-19-EFEMP1^R345W/R345W^ cells make abnormal ECM in vitro.

Our previous experiments using primary RPE cells from *Efemp1*^R345W/R345W^ mice demonstrated that the p.R345W mutation in EFEMP1 causes abnormalities in the structure and turnover of the ECM as well as to the activation of complement *in vitro* [29]. We hypothesized that edited ARPE-19-*EFEMP1*^R345W/R345W^ cells can make abnormal ECM that is similar to the altered BrM found in AMD patients and that precede basal deposits and drusen. We cultured ARPE-19 wildtype and ARPE-19-*EFEMP1*^R345W/R345W^ mutant cells on transwells in RPE media in the absence of serum [29, 40]. After 4 weeks, decellularization of the transwells was performed and the exposed ECM was fixed and imaged by scanning electron microscopy (SEM). SEM images showed that the structure of the ECM generated by the mutant cells was abnormal, with elongated fibers that covered most of the transwell, while the ECM generated by the wildtype cells was organized in a distinct pattern (Fig. 2A-D).

**Figure 2.**
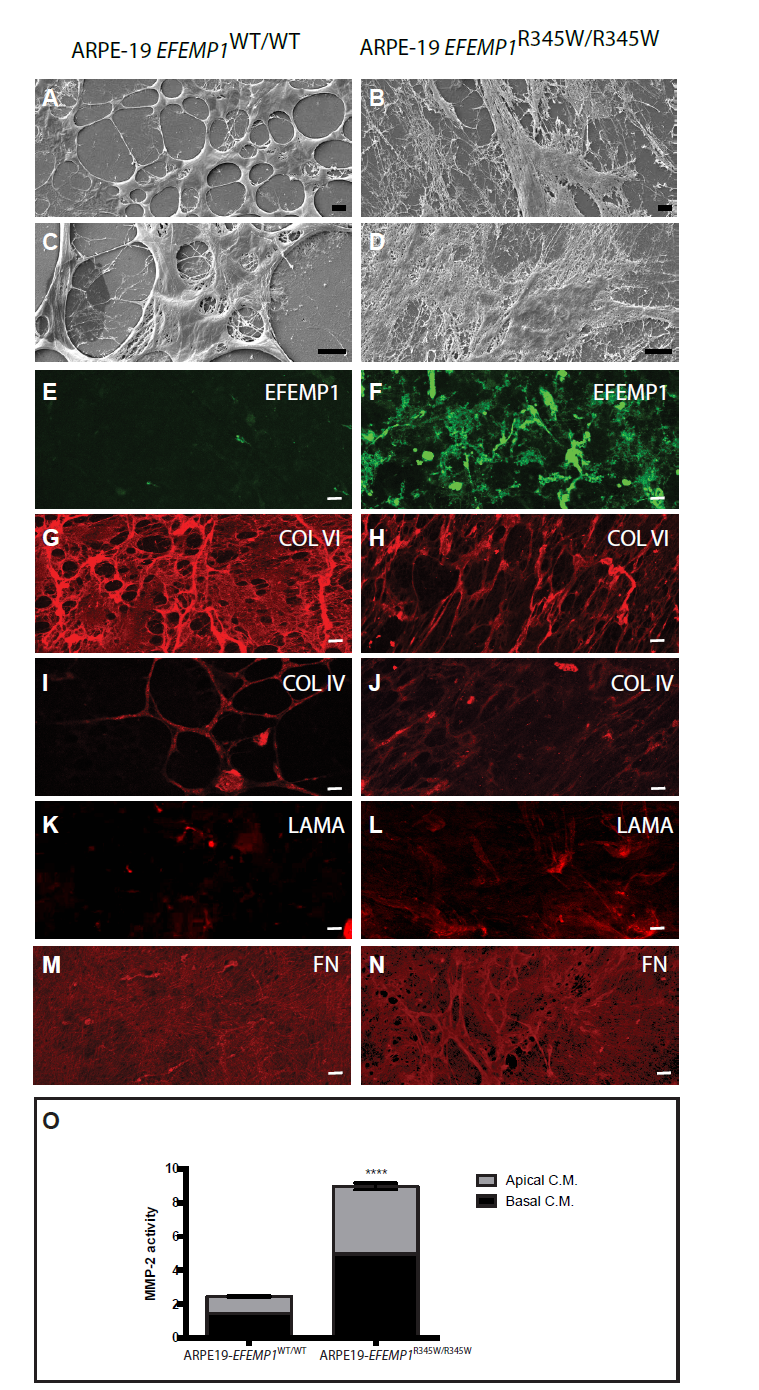
Decellularized transwells of ARPE-19 wildtype (left panels) and ARPE-19-*EFEMP1*^R345/R345W^ (right panels) cultures after 4 weeks. SEM images show normal ECM **(A, C)** vs. abnormal ECM **(B, D)** made by wildtype and mutant cells respectively. Immunostaining with antibodies for EFEMP1 **(E, F),** Col VI **(G, H)**, Col IV **(I, J)**, laminin **(K, L)**, and fibronectin **(M, N)** show the abnormal structure of the ECM. Scale bars A-N: 10 m. **(O)** Relative MMP-2 activity measured by zymography in apical and basal conditioned media of ARPE-19-*EFEMP1*WT/WT and ARPE-19-*EFEMP1*^R345/R345W^ cells. (Data represented as mean ±SD. t-test, ****p<0.0001).

Immunostaining for the main components of basal laminar deposits in AMD patients, such as EFEMP1, laminin, fibronectin, collagen IV and VI [5, 41, 42], confirmed that the abnormal ECM was comprised of these proteins (Fig. 2). The mutant EFEMP1 protein appeared to form extracellular aggregates that extend along the abnormal ECM made by ARPE-19-*EFEMP1*^R345W/R345W^ (Fig. 2E), similar to what we found in primary RPE from *Efemp1*^R345W/R345W^ mutant mice [29]. In contrast, EFEMP1 is barely visible in the ECM made by wildtype ARPE-19 cells (Fig. 2F). Collagen VI labeling in the wildtype ECM is visualized as a continuous network of fibers with some thickened and localized areas (Fig. 2G), while the mutant ECM presents a disorganized network of stretched fibers (Fig. 2H). Collagen IV labeling shows a similar pattern of that collagen VI, with less intense staining (Fig. 2I, J). Weak laminin staining appears to localize in areas where ECM fibers converge (Fig. 2K), and is more extended in the mutant ECM (Fig. 2L). Fibronectin labeling (Fig. 2M) reveals an open network of fibers in the wildtype ECM, while in the mutant ECM the fibronectin network appears to be thicker, with more than one layer of fibers (Fig. 2N).

Zymography analyses demonstrated that MMP-2 activity was significantly increased in conditioned media from ARPE-19-*EFEMP1*^R345W/R345W^ cells compared to wildtype (t-test, p<0.0001), which indicates that the ECM turnover is also affected by the p.R345W mutation (Fig. 2O).

### 3. ECM made by ARPE-19-EFEMP1^R345W/R345W^ cells binds active complement components.

The presence of active complement components anchored to the ECM made by ARPE-19-*EFEMP1*WT/WT and ARPE-19-*EFEMP1*^R345W/R345W^ cells was demonstrated by immunostaining of the exposed ECM after decellularization. Increased deposition of C3b on the abnormal ECM was demonstrated with an antibody that binds C3b once it has been cleaved; but it does not recognize full length C3 (ANOVA, p=0.0172) (Fig. 3). This antibody also binds C3(H2O) generated by hydrolysis of C3 via tick-over. Co-immunostaining with CFH antibodies showed that both C3b/C3(H2O) and CFH are deposited on the abnormal ECM and overlay most of the surface of the abnormal ECM, compared to the wildtype ECM,where the protein deposition is discreet (Fig 3. E, F). We did not detect C3a in conditioned media from edited ARPE-19 wildtype or ARPE-19-*EFEMP1*^R345W/R345W^ cells.

**Figure 3.**
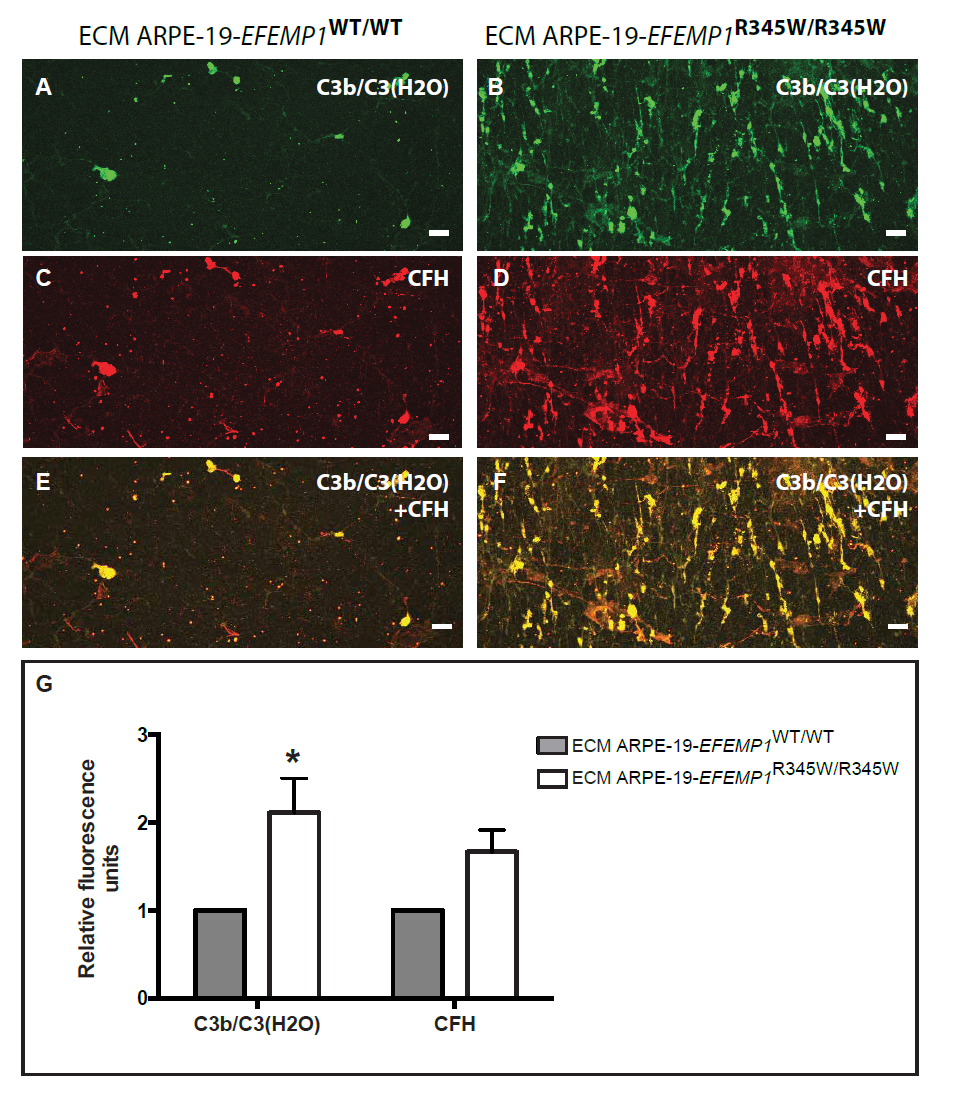
Decellularized transwells of ARPE-19 wildtype (left panels) and ARPE-19-*EFEMP1*^R345/R345W^ (right panels) cultures. Exposed ECM immunostained with antibodies for **(A, B)** C3b/C3(H2O) and **(C,D)** CFH. **(E, F)** Co-staining for C3b/C3(H2O) and CFH. Scale bars 50 m. **(G)** Average quantification of C3b/C3(H2O) and CFH fluorescent labeling comparing exposed ECMs. (Data represented as mean ± SD.*p<0.05).

### 4. Normal primary RPE cells grown on abnormal ECM make basal deposits similar to basal laminar deposits in AMD patients.

To determine if the abnormal ECM made by the ARPE-19-*EFEMP1*^R345W/R345W^ cells can affect the viability and function of normal RPE cells, we isolated primary hfRPE cells and seeded them on the exposed ECM made by ARPE-19-*EFEMP1*WT/WT and ARPE-19-*EFEMP1*^R345W/R345W^ cells on transwells. hfRPE cells were cultured for 2 weeks in the absence of serum. Although the morphology of the cells in both cultures was similar (Fig. 4A, B), transepithelial electrical resistance (TER) after 2 weeks was slightly but significantly lower in cultures grown on abnormal ECM compared to wildtype (t-test, p=0.0123) (Fig. 4C).

**Figure 4.**
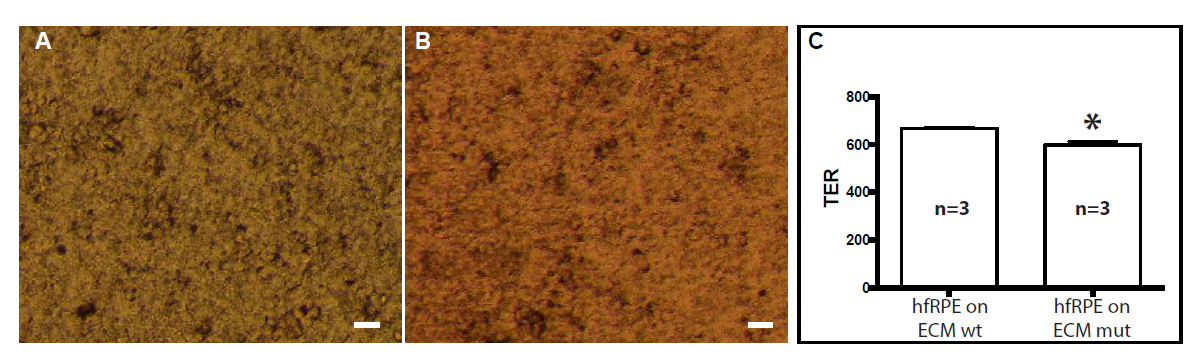
Brightfield microscopy of hfRPE cells grown for 2 weeks on ECM made by **(A)** ARPE-19 wildtype or **(B)** ARPE-19-*EFEMP1*^R345/R345W^ cells. Scale bars 10 m. **(C)** TER measured in hfRPE cell cultured on ECM made by ARPE-19 wildtype or ARPE-19-*EFEMP1*^R345/R345W^ cells for 2 weeks. (n=3 cultures/ECM type. *p<0.05).

To study the formation of basal deposits by normal hfRPE cells grown on abnormal ECM, transwells were decellularized after 2 weeks of hfRPE culture, and the exposed ECM was fixed for immunostaining and SEM. SEM images demonstrated that hfRPE cells grown on normal ECM make typical ECM that covers the insert uniformly (Fig. 5A, C). In contrast, the same hfRPE cells grown on abnormal ECM make thick basal deposits that coat the surface of the transwell as a network of fibers with numerous thickened areas over the original abnormal ECM (Fig. 5 B, D).

**Figure 5.**
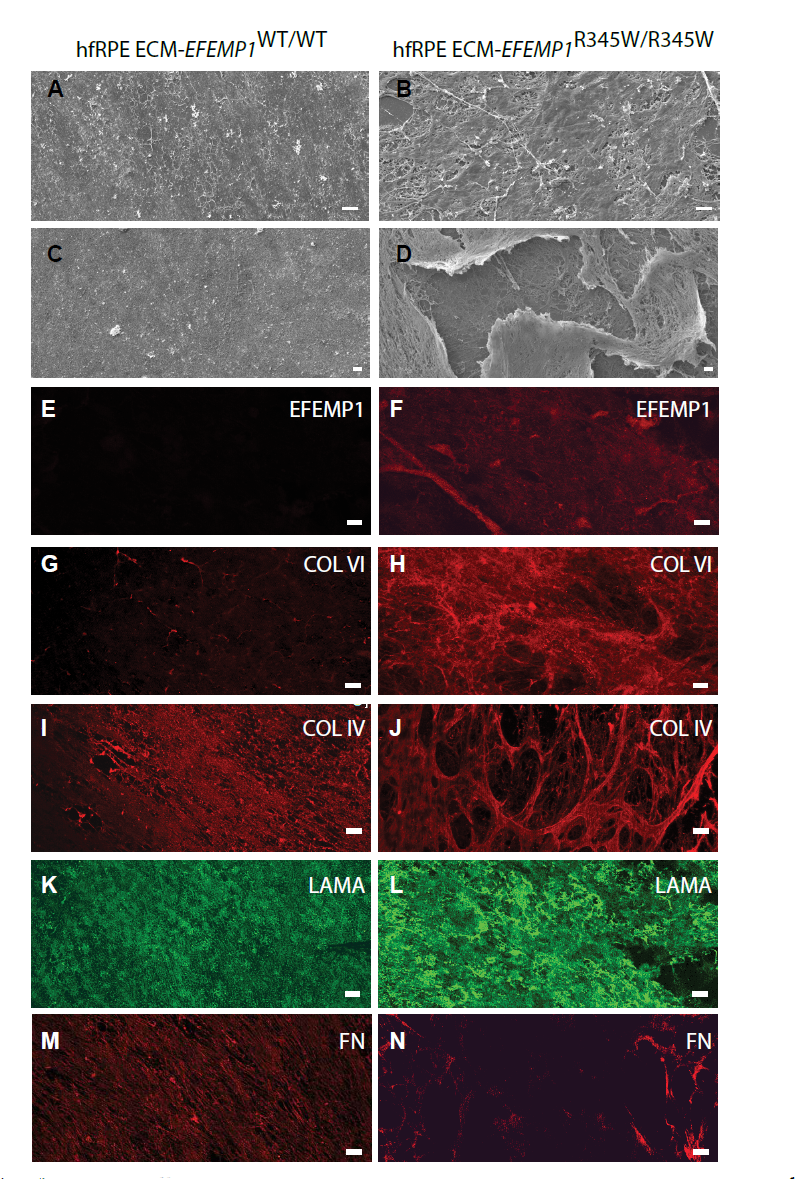
**(A, C)** HfRPE cells grown for 2 weeks on ECM made by ARPE-19 wildtype cells were decellularized and imaged by SEM. Images show the remaining exposed normal ECM made by hfRPE cells. **(B, D)** HfRPE cells grown for 2 weeks on ECM made by ARPE-19-*EFEMP1*^R345/R345W^ cells were decellularized and imaged by SEM. Images show the remaining thick basal deposits made by hfRPE cells. In some of these areas, the outer layer of the deposits is lifted and a network of crosslinking ECM fibers can be observed underneath. Immunostaining of the exposed ECM and deposits with antibodies for EFEMP1 **(E, F)**, Col VI **(G, H)**, Col IV **(I, J)**, laminin **(K, L)**, and fibronectin **(M, N).** Immunolabelling of deposits revealed a very strong signal for collagen VI, and the distribution of both collagen VI and IV was heterogeneous and disposed as a tridimensional network of crosslinked bundles of collagen fibers.The data show that the composition of the deposits made by hfRPE grown on abnormal ECM is similar to the composition basal deposits in AMD patients. Scale bars A-D: 10 μm, E-N: 50 μm

To test if the deposits made by the hfRPE cells were similar to the basal laminar deposits found in AMD patients, immunostaining with antibodies for the main components of the latter (EFEMP1, collagen VI, collagen IV, laminin, and fibronectin) was performed on the exposed ECM and deposits (Fig 5).EFEMP1 was not detected in normal ECM (Fig. 5E), but was clearly present in the basal deposits (Fig. 5F). While normal ECM showed a discreet labeling for collagen VI (Fig. 5G), collagen VI is a major component of the abnormal ECM (Fig. 5H). Collagen IV is present in both the normal and abnormal ECM, although with different patterns of staining (Fig. 5I, J). Laminin had diffuse labeling in both normal ECM and deposits, with apparently stronger staining in the deposits (Fig. 5K, L). Less fibronectin labeling was detected in the deposits with a disrupted staining pattern compared to normal ECM (Fig.5M, N).

### 5. Impaired ECM turnover in RPE cultures with basal deposits.

Aging is associated with thickening of BrM, which is thought to be due to impaired regulation of ECM synthesis and turnover [3-7]. Turnover of BrM, specially the degradation of collagen IV, is controlled by MMP-2 produced by the RPE [8]. To evaluate the impact of alterations in ECM turnover in the production of abnormal ECM by hfRPE, we measured MMP-2 activity in the apical and basal media of hfRPE cells grown on normal and abnormal ECM, produced by ARPE-19-*EFEMP1*WT/WT and ARPE-19-*EFEMP1*^R345W/R345W^ cells, respectively. Increased MMP-2 activity was demonstrated in basal but not apical conditioned media of hfRPE cells grown on abnormal ECM (Fig. 6A) (ANOVA, p=0.0086). The higher levels of MMP-2 only in basal media appear to be a consequence of posttranslational activation of MMP-2 by the abnormal ECM underneath the cells, since the level of MMP-2 mRNA did not change between conditions (t-test, p=0.1769) (Fig. 6B).

**Figure 6.**
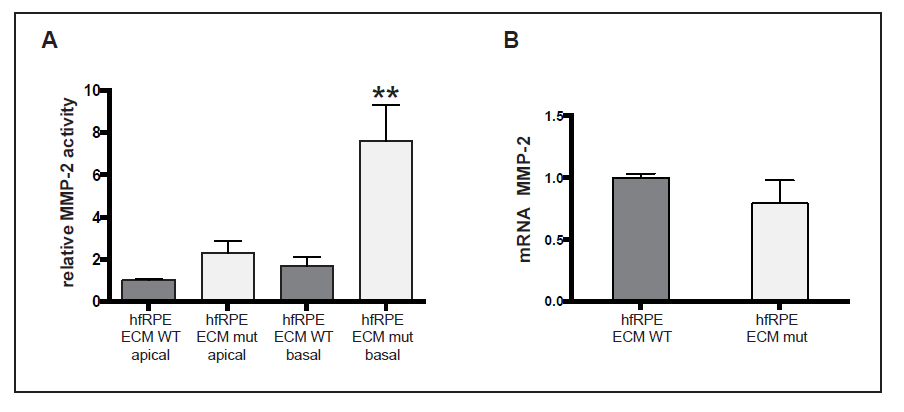
**(A)** MMP-2 activity measured by zymography in apical and basal conditioned media of hfRPE cells cultured on normal (wt) or abnormal (mut) ECM made by ARPE-19-*EFEMP1*^R345/R345W^. **(B)** mRNA levels of MMP-2 measured in the same cultures, normalized to GAPDH. (Data represented as mean ± SEM. **p<0.01. n=4/ECM type).

### 6. Abnormal ECM triggers local activation of the alternative complement pathway by normal RPE cells via tick-over.

We next asked if the abnormal basal laminar deposit material produced by hfRPE cells grown on abnormal ECM was associated with increased local activation of complement via the tick-over process, resulting in increased C3 deposition on the abnormal ECM. Immunostaining of the decellularized ECM generated by hfRPE grown for 2 weeks on normal (made by ARPE-19-*EFEMP1*WT/WT) vs. abnormal (made by ARPE-19-*EFEMP1*^R345W/R345W^) ECM in the absence of serum with antibodies for C3b/C3(H2O) demonstrated increased deposition of activated C3 on the abnormal basal deposit material compared to the normal ECM (t-test, p=0.0279) (Fig. 7A, B). CFH was also detected in areas of strong staining for C3b/C3(H2O) (Fig. 7C-F).

**Figure 7.**
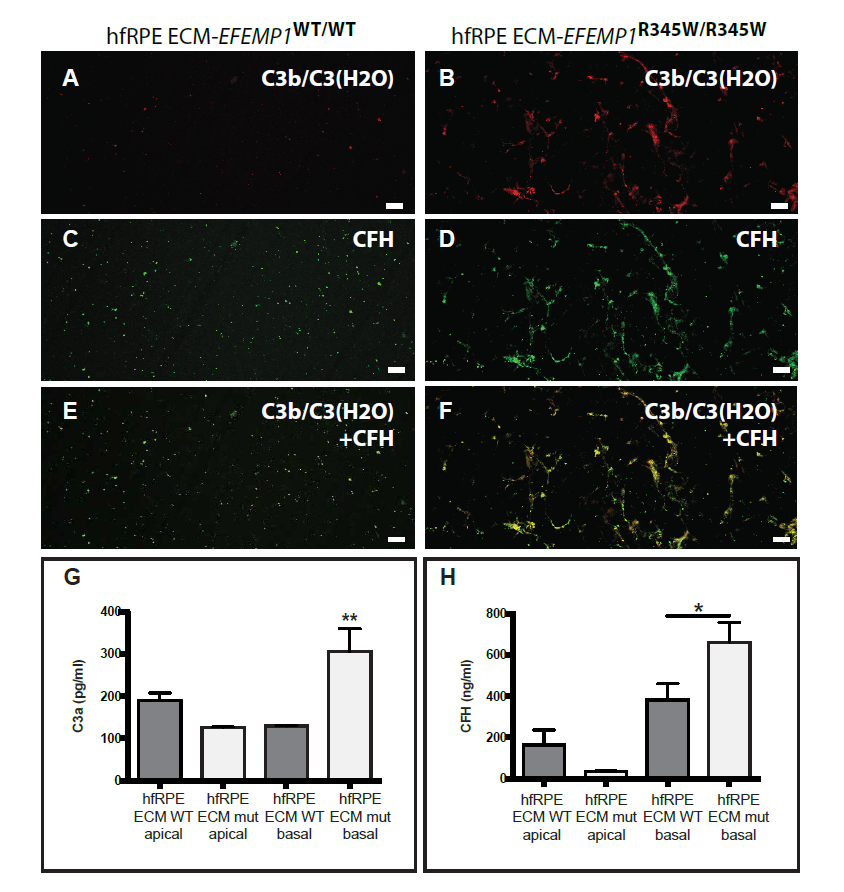
HfRPE cultures grown on ECM made by ARPE-19 wildtype or ARPE-19-*EFEMP1*^R345/R345W^ were decellularized and immunostained with antibodies for **(A, B)** C3b/C3(H2O), **(C, D)** CFH or **(E, F)** both C3b/C3(H2O) and CFH. Scale bars: 50 μm. **(G)** C3a and **(H)** CFH measured by ELISA in apical and basal conditioned media of hfRPE cells grown on ECM made by ARPE-19 wildtype (ECM WT) or ARPE-19-*EFEMP1*^R345/R345W^ (ECM mut) in the absence of serum. (Data represented as mean ± SD. n=4 per type, *p<0.005, **p<0.001).

Unlike classical and lectin pathways, which need the presence of microbial pathogens, alternative complement pathway can be activated via tick-over by deposition of C3b on foreign structures. We tested the hypothesis that induction of the tick-over mechanism via C3b deposition can cause chronic activation of the alternative complement pathway by hfRPE grown on the abnormal ECM. For that, we measured complement activation in conditioned media of hfRPE cells grown for 2 weeks on normal vs. abnormal ECM in the absence of serum. ELISA analyses demonstrated a substantial increase of C3a and CFH in basal conditioned media of hfRPE cells grown on abnormal ECM (Fig. 7G, H) (ANOVA, p=0.0096 & p=0.0219 respectively).

### 7. Alterations in BrM of patients with AMD cause abnormal ECM turnover and complement activation by normal RPE cells.

We hypothesize that increased activation of the complement system by the RPE in response to abnormal ECM is a shared mechanism in *EFEMP1*-associated macular degeneration and AMD. To test this hypothesis, we evaluated the responses of normal hfRPE growth on BrM obtained from human eyes with and without AMD. For these studies, we collected human eyes from age matched donors with and without AMD (ages = 85±1 years). Four normal eyes and four eyes with AMD were used. To evaluate the status of BrM in these eyes, BrM-choroid-sclera explants were evaluated by SEM and immunostaining with antibodies to Collagen IV, which is a major component of the basal lamina in BrM [5, 43]. SEM images demonstrated that BrM of AMD patients had an irregular pattern, where collagen fibers form a network with hexagonal mosaic shape and spotted abnormal structures that resemble drusen (Fig. 8A, B). Collagen IV was present in an irregular pattern in the AMD eyes, compared to more homogeneous labeling in control explants (Fig 8C, D).

**Figure 8.**
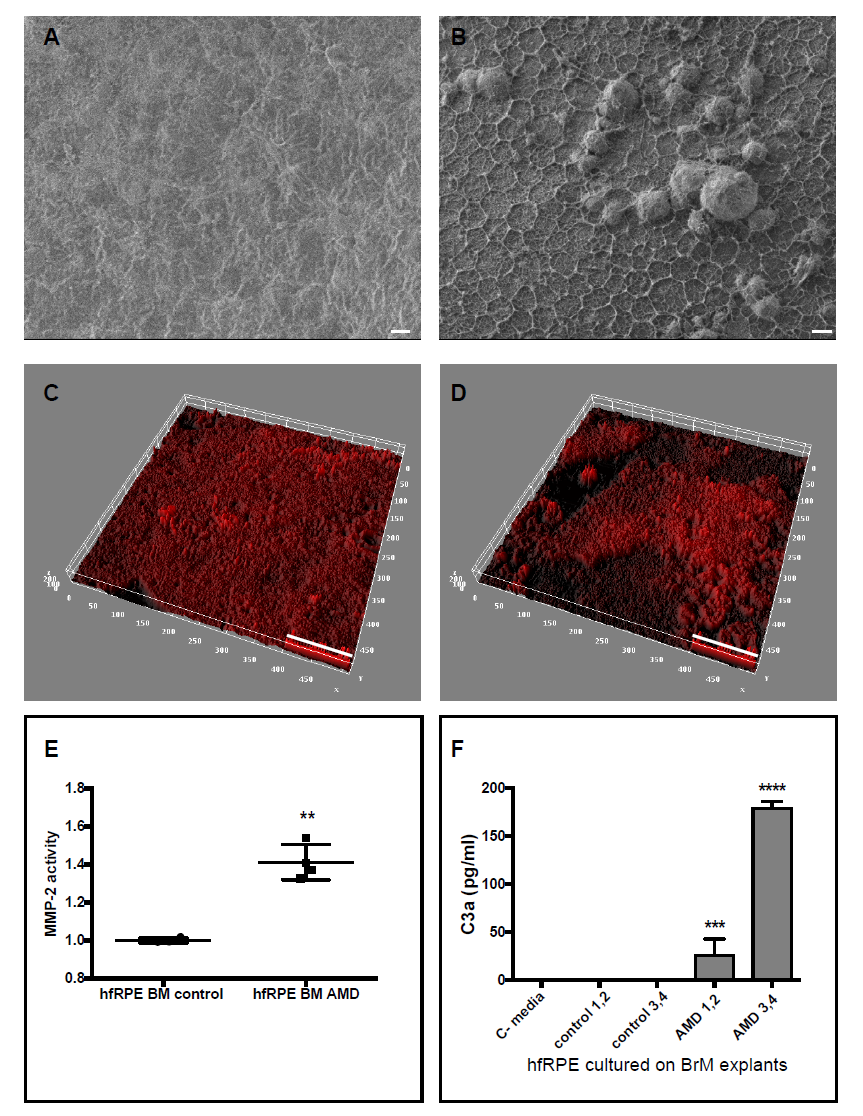
SEM of BrM from **(A)** normal and **(B)** AMD donors. Thick ECM network and drusen-like deposits are visible in the BrM of AMD donors. Representative 3-D reconstruction of BrM explants from**(C)** normal and **(D)** AMD donors immunostained for collagen IV, and imaged by confocal microscope.Quantification of **(C)** MMP-2 activity and **(D)** C3a in conditioned media from hfRPE cells cultured on BrM-choroid-sclera explants from donors without AMD (control 1-4) and with AMD (AMD 1-4). C-media: no cells. (Data represented as mean ± SD. n=4 explants/type. ****p<0.0001,***p<0.001,**p<0.01). Scale bars: (A, B)=10μm, (C, D)=50μm.

We next evaluated the response of hfRPE cells grown on the BrM from the normal and AMD eyes. For these studies, 6mm BrM-choroid-sclera explants were freshly prepared and normal hfRPE cells from the same donor were seeded at a density of 3164 cells/mm2 (approx. 100,000 cells/explant) in RPE medium with 5% FBS. After 48 hours, media was changed and hfRPE cells were cultured on explants in RPE medium for 2 additional weeks in the absence of serum. MMP-2 activity in conditioned media was significantly increased in cultures grown on AMD explants compared to controls, indicating that the ECM turnover is also impaired by the BrM of AMD donors (t-test, p=0.0020) (Fig. 8E). Increased levels of C3a were detected in the conditioned media from hfRPE cells grown on explants with AMD but not media from cells grown on control explants (Fig. 8F) (ANOVA, p<0.0001). These data confirm that the abnormal ECM in the AMD BrM is sufficient to cause local activation of complement system by normal RPE cells.

## DISCUSSION

The results reported here demonstrate that abnormalities in the ECM, including those caused by the p.R345W mutation in EFEMP1 or aging and other risk factors in AMD, are sufficient to cause activation of complement system and formation of basal deposits by normal RPE cells. Like RPE cells from *Efemp1* mutant mice, genome edited ARPE-19-*EFEMP1*^R345W/R345W^ cells make abnormal ECM *in vitro*, which is associated with increased complement activation as indicated by increased levels of C3b in the ECM. The abnormal ECM made by the ARPE-19-*EFEMP1*^R345W/R345W^ cells also stimulated activation of C3 and production of basal deposits by normal hfRPE cells. Of particular note, similar responses were also observed when normal hfRPE cells were cultured on BrM from patients with AMD, suggesting that stimulation of complement activation by abnormal ECM is a shared step in the early pathogenesis of both inherited and age-related macular degenerations. Further, impaired remodeling of the ECM produced by RPE cells also appears to be a shared feature of early disease between inherited and age-related macular degenerations, as increased MMP-2 activity was observed in the basal media of hfRPE cells cultured on the ECM produced by ARPE-19-*EFEMP1*^R345W/R345W^ cells and on BrM from AMD patients. These data provide insight into the early pathogenesis of macular degenerations, and suggest that alterations in complement activation occur earlier in disease than previously described.

Our previous studies using a mouse model of the early-onset *EFEMP1-*associated macular degeneration demonstrated that the formation of sub-RPE deposits *in vitro* occurs in response to local activation of complement system by the RPE [29]. Here, we have shown that abnormal ECM made by ARPE-19-*EFEMP1*^R345W/R345W^ cells has changes in structure and composition similar to those observed in the aged BrM of AMD patients [5, 44]. This abnormal ECM binds active complement components, as demonstrated by increased deposition of C3 and CFH. However, we did not detect C3a in the conditioned media from these cells, perhaps because the concentration was too low to be detected. The abnormal ECM observed may be due to impaired ECM turnover, since increased MMP-2 activity was present in conditioned media from ARPE-19-*EFEMP1*^R345W/R345W^ cells. The increased MMP-2 activity may be due in part to decreased inhibition of MMP activity by TIMP-3, a protein that interacts with EFEMP1 by the domain that contains the R345W mutation [45].

We tested the hypothesis that changes in the structure and composition of BrM in early AMD affect the function of RPE and generate inflammation. Precise genome editing of the human ARPE-19 cell line carrying the *EFEMP1*^R345W/R345W^ mutation allowed developing a model that recapitulates the RPE/BrM pathology at early stage of AMD. SEM and immunostaining analyses demonstrated that normal hfRPE cells grown on abnormal ECM produced by ARPE-19-*EFEMP1*^R345W/R345W^ cells make thick basal deposits with increased collagens VI and IV, in addition to a distinct pattern for ECM components of basal laminar deposits. The abnormal deposition of basement membrane material underneath the RPE is also a characteristic feature of early stage AMD [5, 41, 42, 44, 46, 47]. In addition, explants of BrM with AMD also showed increased deposition of ECM and collagen IV. Accordingly, our model is an effective tool to study the mechanisms underlying the formation of basal deposits, which is a hallmark of early AMD. Moreover, it demonstrates that modeling some aspects of complex diseases using Mendelian diseases is feasible with CRISPR-Cas9 mediated genome editing, the efficiency of which can be improved by the addition of the DNA ligase IV inhibitor, SCR7 [48].

It is currently accepted that MMPs are specific proteolytic enzymes rather than mere ECM degraders. In this way, MMPs are key for the homeostasis of the extracellular environment and regulation of innate immunity [49]. Aberrant ECM expression and remodeling can trigger the activation of immune cells in chronically inflamed tissues [10]. In the case of BrM, MMP-2 produced by the RPE is crucial for the degradation of the basement membrane, which increment with age results in BrM thickening, altered ECM turnover and inflammation [5, 6, 8, 9, 50]. The increased MMP-2 observed in hfRPE cultures grown on abnormal ECM indicates that the ECM turnover and remodeling is impaired, which causes further abnormal deposition of ECM proteins. It is also probable that the RPE secretes MMP-2 in an attempt to recover ECM homeostasis by eliminating the collagen accumulated underneath. Proteolysis of collagen results in bioactive fragments that promote the local accumulation of debris, cytokines, chemokines and other ECM proteolytic fragments and regulators, which could explain the formation of basal deposits [4, 6, 10-12]. The abnormal accumulation of these substances in the ECM along with the abnormal structure of the interface between the RPE and BrM may lead to complement activation. Our data determined that abnormal ECM stimulates the activation of complement by hfRPE cells as shown by increased C3a and CFH in the basal media. HfRPE cells also show increased complement activation, as demonstrated by C3a in media, following growth on BrM from AMD compared to control eyes. Hence, the mechanism of abnormal ECM causing increased complement activation is also shared between *EFEMP1*-associated macular degeneration and AMD.

In contrast to specific interactions that initiate classical and lectin pathway activation by pathogens, the alternative pathway can be autoactivated via tick-over by covalent attachment of C3b to nearby surfaces [51]. Although the tick-over is a basal process, it can be actively induced when C3(H2O) is deposited on abnormal ECM [33-35]. Specifically, C3(H2O) has great affinity to bind collagen IV and laminin [35], which are major components of the abnormal ECM and basal deposits made by the mutant RPE cells in our model. We believe that C3(H2O)-convertase stabilizes on the abnormal ECM, evading CFH inhibition, which results in a chronic activation of the alternative complement pathway and increased CFH and C3a in media [31, 36, 37]. Our hypothesis is supported by the fact that only hfRPE cells seeded on abnormal ECM or BrM with AMD in the absence of pathogens showed increased C3a, while no complement activation was detected when the same cells were grown on normal ECM or BrM without AMD. Taken together, these data suggest that the primary activation of the complement system in the deposits occurs via tick-over and posterior amplification of the alternative pathway. However, additional analyses are needed to corroborate this supposition.

To our knowledge, this is the first demonstration that abnormal matrix can initiate the local activation of complement system as one of the early steps in the pathogenesis of AMD. The process has similarities with other diseases such as fibrosis, arthritis, kidney disease, Alzheimer disease, and cancer, in which the excessive deposition of ECM triggers inflammatory responses [52-56]. However, other factors such as genetic variants that alter the responsiveness of the complement system are needed to initiate disease in AMD [15]. These data help define the earliest stages in macular degeneration disease pathogenesis.They also confirm the importance of complement system in dry AMD, and most importantly, they suggest that alterations in complement activity participate in disease earlier than previously described [13, 14, 57].

In conclusion, the efficacy of therapies for the treatment of early AMD might be improved using regulators of the ECM synthesis and turnover, a target that has shown to be key in other human diseases like cancer, fibrosis, and cardiovascular disease [58-60]. Also, new drugs addressing the specific steps involved in the RPE, such as the tick-over process, may be efficient at early stages of AMD. Moreover, a combination of conventional therapy, such as complement-modulating drugs, with the ECM regulators applied locally could be an improved approach for patients with drusen, and could stop or at least delay the progression of the disease to legal blindness without compromising the whole immune system or vital biological processes. Further, our model can be used to test different treatments *in vitro*, as well as to better understand the mechanisms that lead from BrM thickening to basal deposit formation and disease progression in AMD.

## METHODS

### CRISPR/Cas9 genome editing

#### Single guide RNA (sgRNA) design

ARPE-19 cells (ATCC® CRL-2302™, Manassas, VA) were edited using the Clustered Regularly Interspaced Short Palindromic Repeats (CRISPR) technology as previously described [38, 39]. The single guide sgRNA target sequence (GACCACAAATGAATGCCGGG) was designed with the tool http://crispr.mit.edu/, with a score of 82. All potential off-targets have at least 2 mismatches and a maximum score of 2.2. Potential off-targets with a score >0.2 were ruled out by PCR followed by Sanger Sequencing. The sgRNA was cloned onto the vector pSpCas9(BB)-2A-GFP (PX458) (a gift from Feng Zhang, Addgene plasmid # 48138) using the BbsI site to be expressed under the U6 promoter.

#### Transfection of ARPE-19 cells

ARPE-19 cells were transfected with the Amaxa nucleofector kit V (Lonza, Portsmouth, NH) following manufacturer’s instructions. 5µg of plasmid DNA was co-transfected with 5µl of 10µM ssODN donor (5’ T CTC TGG TGT TAG AAT GTA GGG ATC TTG ACA AGG ATT TCG TGG ATA ACA ACG GAA GCC GCC ATG ATA ATT CCA ACA CAT TTC ATC TTC CCA GCA TTC ATT TGT GGT CTC ACA CTC ATT TAT GTC CGT AGA TAT GTA GGG TCA AAG AGT TTA CTA ACT AAA CTA ATG AAC TGA TCT AAT TAA 3’) per 10^6 cells in a 10cm dish. Silent mutation was introduced to the PAM sequence in order to avoid cuts in the ssODN (Fig. 1). After transfection, the cells were cultured in DMEM:F12+10% FBS in the presence of 1µM of SCR7 (ApexBio, Houston, TX), a DNA ligase IV inhibitor [48, 61], for 48 hours.

#### Cutting efficiency

was tested using the SURVEYOR assay 48h post-transfection as previously described [39]. Briefly, cells were lysed and DNA was extracted using 10µl of the QuickExtract DNA extraction solution (Epicentre, Madison, WI) per 96-well, and 1µl was amplified using the primers F: 5’ TCCCCCTGGCAAAATTACCC 3’ and R: 5’ AGTTGTGGCCTGTATCTGGA 3’ following the conditions published by Ran et al. [39]. 400ng of PCR product were used to form the heteroduplex, later digested with 2ud of T7 Endonuclease I (New England Biolabs, Ipswich, MA) for 30 min at 37°C. Fragments were resolved in a 2.5% agarose gel.

#### Isolation of clonal cells

was performed by limit dilution as previously described [39]. Although the vector pSpCas9(BB)-2A-GFP (PX458) has GFP and it can be sorted by fluorescence, pilot experiments showed that recovery efficiency for ARPE-19 was very low after sorting compared to limit dilution.Thus, the GFP signal was only used to estimate the rate of transfection under fluorescent microscope (>90%). 48h post-transfection cells were diluted to 1cell/well in 96-well plates and expanded until 60% confluence was reached (around 4 weeks). Then 2 replica plates were prepared, one to keep cells growing in incubator and one to analyze the genotype of the edited clones [39]. After one week, wells with no clones or more than one clone were discarded.

#### Genotype of edited clones by PCR-RFLP

DNA was extracted from replica plate using the QuickExtract DNA extraction solution (Epicentre, Madison, WI) as previously described [39]. The change C>T c.1034 in the *EFEMP1* gene creates a new restriction site for the *BseYI* enzyme, which allows for the rapid screening of positive edited clones by PCR-RFLP before Sanger sequencing (Fig. 1B). PCR fragments amplified using the primers F: 5’ TGTCCCTGATGACAAGAACTGG3’ and R: 5’ CTCGGCACATGGCATTTGAG 3’ (amplicon size 569bp) were digested with *BseYI*, generating 2 fragments of 364bp and 215bp when the recombination had taken place. The positive C>T c.1034 knockin clones were sequenced by Sanger to rule out unwanted indels and off-targets. The counterpart replica wells maintained in culture were expanded in DMEM:F12+10% FBS and used for subsequent experiments or cryopreserved.

Hemizygous samples were genotyped by cloning above-mentioned PCR fragments with the TA Cloning® Kit, with pCR™2.1 Vector and One Shot® TOP10 Chemically Competent *E. coli* (Invitrogen/Thermo Fisher Sci., Waltham, MA) following the manufacturer’s instructions. Minipreps were performed using Zyppy™ Plasmid Miniprep Kit (Zymo Research, Irvine, CA) and sequenced by SANGER.

### Generation of ECM by ARPE-19-*EFEMP1*WT/WT and ARPE-19-*EFEMP1*^R345W/R345W^ cells

Selected clones of ARPE-19 cells carrying none or 2 copies of the mutation C>T c.1034 were seeded on 12-mm polyester transwells (Corning, NY) at a density of 4x10^4 cells per transwell in DMEM:F12+10%FBS per triplicate. After 4 weeks, transwells were decellularized by incubating them with sterile 0.5% Triton X-100 + 20mM NH4OH in PBS (1.5ml was added to the bottom chamber and 0.5ml to the apical chamber) for 5 min at 37&3x00B0;C, followed by several washes with sterile PBS. hfRPE cells were seeded immediately on the exposed ECM or inserts were fixed for subsequent analyses.

### HfRPE isolation and culture

Eyes from 16-20 weeks of gestation fetuses were obtained from Advanced Bioscience Resources (Alameda, CA) placed in RPMI on ice, and delivered by an overnight priority delivery service. All tissues were used less than 24 hours after enucleation. Primary hfRPE cells were collected as previously described with minor modifications [40, 62]. Briefly, upon reception eyes were rinsed in antibiotic-antimycotic solution for 5 min at RT. Connective tissue was removed and the anterior portion of the eye, cornea, iris, and vitreous were also removed. Eyecups were incubated in Dispase I solution (2U/ml in RPE media+5%FBS) for 25 min at 37°C. Neural retina was then removed and the RPE sheet was peeled off with fine forceps and incubated in 0.25%Trypsin+ 0.02% EDTA for 10 min at 37°C in a water bath. After centrifugation, cells from one eye were resuspended in RPE media+15%FBS and seeded on a total of 6 transwells, 3 decellularized transwells containing ECM made by ARPE-19-*EFEMP1*WT/WT and 3 containing ECM made by ARPE-19-*EFEMP1*^R345W/R345W^. Confluence after 24 hours was around 30%. FBS was removed from the media after 72 hours. After 2 weeks, transwells were decellularized as previously described and fixed in 4% PFA for subsequent analyses.

**\*RPE media** was pre pared as previously described [40, 63]. N1 Medium Supplement 1/100 vol/vol, glutamine 1/100 vol/vol, penicillin-streptomycin 1/100 vol/vol, and nonessential amino acid solution 1/100 vol/vol, hydrocortisone (20 µg/L), taurine (250 mg/L), and triiodo-thyronin (0.013 µg/L) in alpha MEM [40] + 5% heat inactivated fetal bovine serum (FBS) (Hyclone, Logan, UT) or without FBS.

**Transepithelial electrical resistance (TER)** of the RPE cultures were measured using an epithelial voltohmmeter (EVOM) [64] [40].

### Preparation of BrM explants

BrM explants were obtained from human donors through the National Disease Research Interchange (Philadelphia, PA). Eyes from 2 donors diagnosed with AMD and 2 normal donors (ages 85±1) were enucleated within 10 hours postmortem and shipped to our facility within 24 hours in a sterile container in DMEM on ice. BrM explants were prepared upon receipt as previously described [65]. Briefly, eyes were washed in 10% povidone iodine for 10 min two times at RT, then rinsed with DMEM containing antibiotics. A circular incision was made below the ora serrata to remove cornea, lens, iris, vitreous and neural retina. RPE cells were removed by incubation with sterile 20mM NH4OH+0.5% Triton in PBS for 20’ at 37C. RPE cells were then flowed out with PBS 1X using a p1000 micropipette. Eyecups were inspected under dissection microscope to ensure the complete removal of RPE cells. 6-mm explants containing BrM-choroid-sclera were performed with a trephine, and placed onto 96-well plates in PBS. Explants were fixed in 4% PFA for 10 min for subsequent analyses by immunostaining and SEM, or used fresh to seed hfRPE cells at a density of 3164 cells/mm^2^ (approx. 100,000 cells/explant). Cells were grown in RPE media with 5% of FBS for 48 hours and then in the absence of serum for 2 additional weeks with 3 changes of media.

### Immunostaining

Transwell inserts containing exposed ECM after decellularization of cultures were rinsed in PBS, fixed for 10 min in 4% paraformaldehyde (PFA) in PBS followed by fixation in 1% glutaraldehyde for 30 min at room temperature. Then inserts were cut off with a razor blade and stored upside in PBS at 4°C pending immunohistochemical analyses. Inserts were cut into small pieces and blocked with 1% BSA for 30 min at RT. Primary antibodies were incubated overnight at 4°C. Secondary antibodies labeled with Alexa-488 or Alexa-555 (Life Technologies, Grand Island, NY) were incubated for 1 h at RT. Sections were mounted with Fluoromount G (Electron Microscopy Sciences, Hatfield, PA) and visualized by TCS SP5 II confocal laser scanning microscope (Leica). Controls were incubated only with secondary antibody. Primary antibodies used were: Col IV (AB6586, Abcam, Cambridge, MA), Col VI (AB6588), LAM (AB11575), FN (AB2413), EFEMP1 (SC33722, Santa Cruz Biotechnology, Santa Cruz, CA) C3b/C3(H2O) (HM2286, Hycult Biotech, Plymouth Meeting, PA), CFH (AB53800).

For the immunostaining of BrM, explants of BrM-choroid-sclera were incubated with antibodies as described above in 96-well plates. After incubation with the secondary antibody, explants were incubated with 0.1% Sudan Black B in 70% ethanol for 5 min at RT to remove autofluorescence, and then washed with 70% ethanol to eliminate the excess of dye. BrM-choroid was then peeled off and flat mounted (oriented with the BrM side up) with Fluoromount G on a slide. Samples were visualized by TCS SP5 II confocal laser scanning microscope (Leica).

### Quantification of the fluorescent signal

Images were converted to binary format with ImageJ [66]. The integrated intensity was measured.

### Tridimensional reconstruction of BrM explants

z-stack images were taken every 0.5 microns, then 3-D surface was reconstructed and plotted using ImageJ [66].

### SEM

Transwell inserts containing exposed ECM after decellularization of cultures and BrM explants were fixed as previously described. Samples were washed in PBS, hydrated in dH2O for 5 min and dehydrated by serial ethanols, 35%, 50%, 70%, 95%, 95% and 100% followed by critical dehydration using the SAMDRI-795 system [63]. After dehydration, specimens were coated with Chromium using a Gatan Ion Beam Coater for 10 min and imaged by Field Emission Scanning Electron Microscope (JEOL 7401F).

### ELISA

Conditioned media from cells cultured on transwells or on BrM-choroid-sclera explants were collected after 2 weeks and concentrated to equal volumes through 3 kDa Amicon filters (Millipore, Billerica, MA). The fraction over 3 kDa was used to quantify C3a and CFH using ELISA kits from R&D Systems (Minneapolis, MN) following manufacturer’s instructions. All samples were assayed at least per duplicate.

### Zymography

10 µl of the above concentrated media were loaded onto Novex 10% gelatin gels (Life Technologies, Grand Island, NY). Zymography assays were then performed per manufacturer’s instructions. Gels were scanned using the Odyssey system (Li-Cor, Lincoln, NE). Gelatinase activity was quantified through the intensity of the bands with the software ImageStudioLite (Li-Cor, Lincoln, NE).

### Statistical analyses

Results are expressed as mean ±SD, with p<0.05 considered statistically significant. Differences between groups were compared using the Student t-test or ANOVA as appropriate using the GraphPad Prism software.

## ACKNOWLEDGEMENTS

The authors would like to thank Ann Tisdale for helpful technical assistance with SEM.

